# Rapid and collective determination of the complete “hot-spring frog” mitochondrial genome containing long repeat regions using Nanopore sequencing

**DOI:** 10.1101/2022.12.21.521512

**Authors:** Yuka Asaeda, Kento Shiraga, Makoto Suzuki, Yoshihiro Sambongi, Hajime Ogino, Takeshi Igawa

**Affiliations:** Graduate School of Integrated Sciences for Life, Hiroshima University; Amphibian Research Center, Hiroshima University

## Abstract

The mitochondrial genome (mt-genome) is one of the promising molecular markers for phylogenetics and population genetics. Recently, various mt-genomes have been determined rapidly by using massively parallel sequencers. However, the control region (CR, also called D-loop) in mt-genomes remain difficult to precisely determine due to the presence of repeat regions. Here, using Nanopore sequencing, we succeeded in rapid and collective determination of complete mt-genome of the hot-spring frog, *Buergeria japonica*, and found that its mt-genome size was 22,274 bp including CR (6,929 bp) with two types of tandem repeat motifs forming repeat regions. Comparison of assemble strategies revealed that the long- and short-read data combined together enabled efficient determination of the CR, but the short-read data alone did not. The *B. japonica* CR was longer than that of a congenic species inhabiting cooler climate areas, *Buergeria buergeri*, because of the long repeat regions in the former. During the thermal adaptation of *B. japonica*, the longer repeat regions in its CR may have accumulated within a short period after divergence from *B. buergeri*.

## Introduction

Recent advances in massively parallel sequencing technologies have facilitated the rapid acquisition of genomic information, leading to the development of a variety of applications. Such technologies also apply to mitochondrial genomes (mt-genomes) [e.g., 1–3], which are used as molecular markers for phylogenetics and population genetics. However, the control region (CR, also called D-loop) in the mt-genome of some animals is difficult to sequence due to the presence of repeat sequences in such species. To overcome this problem, long-read sequencing technologies [4,5] are beginning to contribute to the determination of sequences that are to-date difficult to read.

In amphibians, the CR includes direct and inverted tandem repeat sequences causing longer total length of their mt-genome. Moreover, frequent gene rearrangements have also been reported in modern anuran amphibians (Suborder Neobatrachia) [6]. The mt-genome of a bell-ring frog, *Buergeria buergeri*, also includes long CR of approximately 4.6 kbp and rearranged ND5 gene (translocation of ND5 next to CR) [7]. However, the evolutionary origin of these mt-genome features has not been clarified because only one species of genus *Buergeria* has been reported so far. Recent molecular phylogenetic analyses based on several mitochondrial genes reveal that *B. japonica*, inhabiting widely across Taiwan and the Ryukyu Archipelago in Japan, is the first diverged species within genus *Buergeria* [8,9]. Thus, sequencing the *B. japonica* mt-genome can reveal the origin of mt-genome features in this genus. Apart from being important molecular markers, mitochondrial genes, which are involved in aerobic energy metabolism, are also known to contribute to the organismal temperature tolerance function in some organisms [10,11]. Because *B. japonica* is also known as “hot-spring frog” and the tadpoles have high temperature tolerance [12–15], clues to understanding the evolutionary strategies for temperature adaptation will be available with the determination of the *B. japonica* mt-genome sequence, in comparison with that of *B. buergeri* inhabiting cooler climate areas. In the present study, we conducted sequencing and assembling of the *B. japonica* mt-genome. We first obtained sequence data by Illumina and Oxford Nanopore Technology (ONT) sequencers, and then examined multiple recent assembly software. The results demonstrate the successful use of sequencing and assembly strategies to obtain the complete *B. japonica* mt-genome, which can then help to shed light on organismal thermal adaptation in this species.

## Materials and Methods

### Extraction of genomic DNA

A female *B. japonica* individual was collected in April 2018 at Seranma hot spring in Kuchinoshima, Tokara Islands, Japan, and kept until death in March 2019 at the Amphibian Research Center, Hiroshima University. Genomic DNA was extracted from the muscle of fresh dead body of this individual using DNA suisui-F (Rizo, Tsukuba, Japan) according to the manufacturer instructions and dissolved in 50 μL of nuclease-free water.

### Genomic DNA sequencing using Illumina sequencer

The extracted genomic DNA was used for library construction using NEBNext® DNA Library Prep Kit (New England Biolabs, MA, USA), and sequencing (150 bp paired end) was conducted at Novogene Co. Ltd. (Beijing, China) using Novaseq 6000 sequencing.

### Genomic DNA and amplicon sequencing using Nanopore sequencer

The extracted genomic DNA was also used for library construction for Nanopore sequencing using a ligation library preparation kit (LQK-LSK109, ONT). Following manufacturer instructions, the quality and molecular weight of the genomic DNA were measured using Qubit, and sequencing was conducted using MinION sequencer (MinION Mk1B, ONT) and Flongle flow cell (FLO-FLG001, ONT).

To perform sequencing more effectively, we also conducted amplicon sequencing. For amplification of mitochondrial DNA fragments by PCR, two sets of oligo DNA primers were designed for two regions (spanning from ND6 to 12S rRNA and 12S rRNA to Cyt*b*) based on the sequence of *B. buergeri* (AB127977.1) [7] and partial sequences of Cyt*b* from *B. japonica* (AB998751-63) [16]: Bb_ND6_13766_Fow 5’-CTCGGACACCCCTCATCACTCA-3’; Bb_12SrRNA_2722_Rev 5’-GAGCTGCACCTTGACCTGACGT-3’; Bb_12SrRNA_2559_Fow 5’-CAACGCCAGGGAATTACGAGCT-3’; Bj_Cytb_Rev 5’-AGGATTTTTGTAAGTGGGCGGAA -3’. Polymerase chain reaction (PCR) amplification was performed in a Bio-Rad Laboratories T100 thermal cycler in 20 µL reactions containing 10 µL KOD One^®^ PCR Master Mix (TOYOBO, Osaka, Japan), 1 µL DNA solution, and 10 pmol of each primer. Temperature cycle was performed in an initial step of 95 °C for 3 min, followed by 35 cycles of 98 °C for 10 s and 68 °C for 3 min. PEG (polyethylene glycol) precipitation [17] was then conducted to remove the remaining primers and the sample was dissolved in 10 μL of nuclease-free water. PCR products (amplicons) were measured for DNA concentration using Qubit Fluorometer (Thermo Fisher Scientific) and Qubit dsDNA HS Assay Kit (Thermo Fisher Scientific), and 150 ng of each amplicon was used for library preparation with the LQK-LSK109 ligation library preparation kit. Library preparation and sequencing were conducted in the same manner as with genomic DNA sequencing. For sequence analysis, we used a PC with NVIDIA Quadro P2200 and MinKNOW software for basecalling (ONT). Real-time basecalling was performed in super accuracy basecalling mode.

### De novo assembly of mt-genome

We attempted assembly of the mt-genome of *B. japonica* using three different datasets: (1) Illumina short read data (genomic DNA, 150PE); (2) Nanopore long read data (genomic DNA); and (3) Nanopore long read data (amplicon) (Fig. 1). For dataset (1), we used MitoZ 2.4a [18] with “all” command trimming of adapter and low-quality sequences using trimmomatic v0.39 [19]. For dataset (2), we used a previously developed pipeline [20]. The reads derived from mt-genome were extracted using mtBlaster (https://github.com/nidafra92/squirrel-project/blob/master/mtblaster.py) after filtering reads with average quality score of 10 or higher using NanoFilt [21]. The mt-genome sequence of *B. buergeri* (AB127977.1) was used as reference data. The extracted sequence data was assembled using minimap2 v0.3-r179 [22] with the runtime option ava-ont and miniasm v2.24-r1122 [22]. The resultant contigs were polished using racon v1.5.0 [23] after re-mapping the raw reads to the contigs using minimap2. To improve sequence accuracy, polishing was performed five times. Finally, the polished sequence was indexed using BWA-mem2 [24] and polished again using Nanopolish (https://github.com/jts/nanopolish). For dataset (3), we assembled reads using Trycycler [25] after filtering reads with an average quality score of 10 or higher and a length of 1,000 bp or greater using NanoFilt. The resulting contigs were filtered by length between 15,000 bp and 25,000 bp using ‘seq’ command of seqkit [26] and aligned and merged in MEGA X [27]. Finally, after comparing the three assemblies, the complete mt-genome sequence of *B. japonica* was determined by combining the coding region based on (1) and the CR based on (3). The length of the CR was verified by checking distribution and frequency of the length in raw reads in (3) using electronic PCR (ePCR) [28] with the 5’ and 3’ tip of the CR as a primer pair. The protein coding genes were annotated based on information of the mt-genome of *B. buergeri* using the ‘Annotate from Database’ function of Geneious (version 9.1.8).

**Fig. 1.**
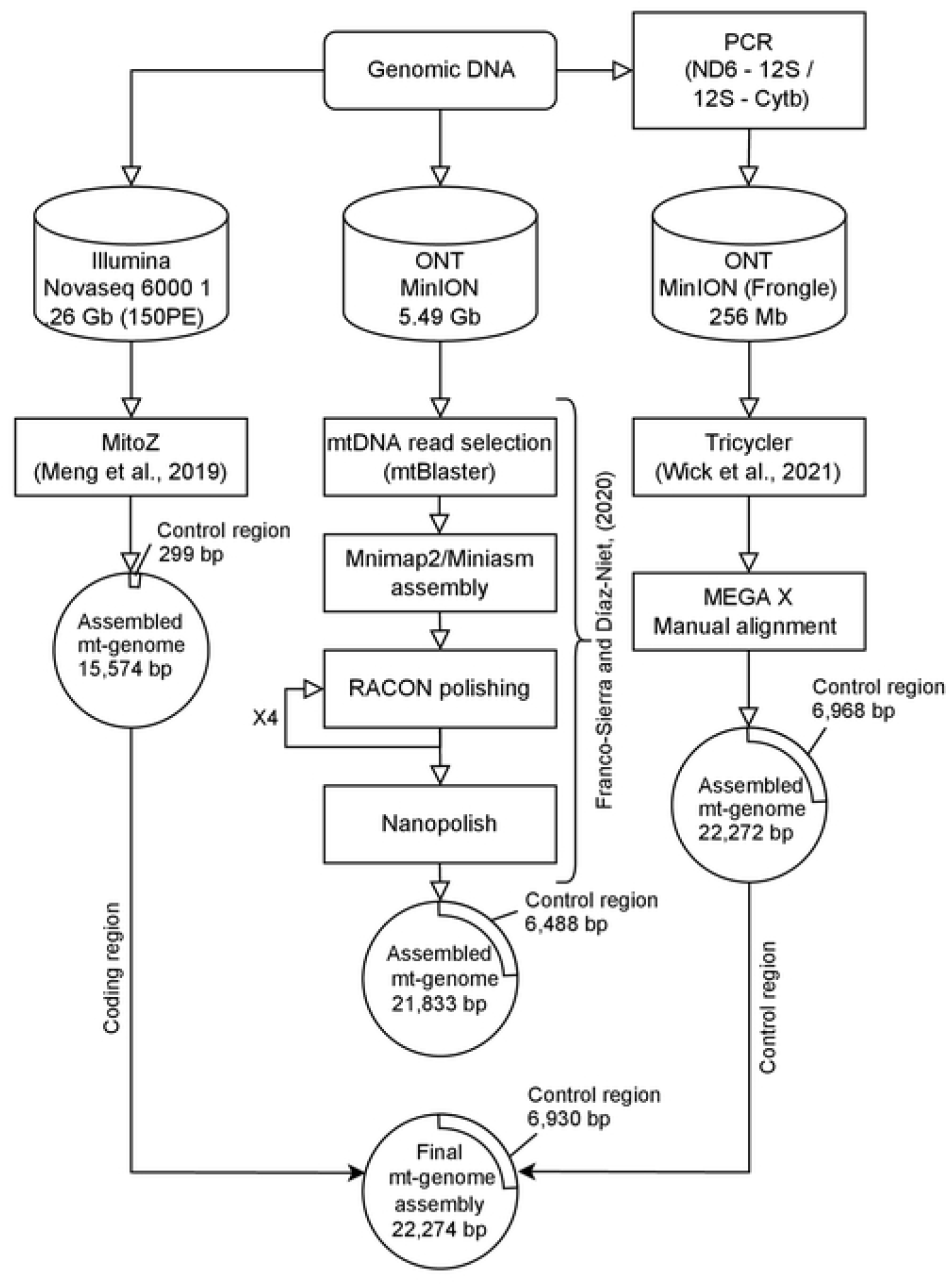
Flowchart of mitochondrial genome assembly of *B. japonica* using Illumina and Nanopore sequence data.

## Results

The total yield of *B. japonica* genomic DNA using Illumina sequencer (1) was 1,256,313,437 bp (8,375,809 paired reads), whereas yields of genomic DNA (2) and PCR amplicon of mt-genome (3) using ONT MinION/Flongle sequencer were 5,491,346,786 bp (total reads: 1,106,685, read length N50: 9,354 bp, read length average: 4,962 bp) and 255,561,618 bp (total reads: 80,109, read length N50: 10,107 bp, read length average: 3,190 bp), respectively (Table 1). Data were deposited in the DDBJ Sequence Read Archive (DRA) under accession numbers DRR418996–8. De novo assembly of the short reads (1) using MitoZ assembler resulted in a *B. japonica* mt-genome size of 15,574 bp with 943 mean coverage depth per nucleotide. In contrast, assembly of the long reads of genomic DNA (2) using the pipeline [20] and of amplicon (3) using Trycycler [25] resulted in mt-genome sizes of 21,833 bp (with 270 mean coverage depth per nucleotide) and 22,272 bp (with 2,253 mean coverage depth per nucleotide), respectively.

The CR sequences obtained in (1), (2), and (3) differed in length: 229 bp, 6,488 bp, and 6,968 bp, respectively. The long-read assemblies in (2) and (3) contained two types of tandem repeat motif sequences [20–24 of repeat motif A (40 bp) and 168–203 of repeat motif B (20 bp)] (Table 1). These repeat motifs were also found in assembly of (1), but with only small copy number (5 for motif A and 1 for motif B). The number of repeat motifs varied even in assembly of (2) and (3), and thus, we measured the length of CR in raw reads in (2) and (3) by ePCR. Histogram of the length of CR exhibited unimodal distribution with 6751–6850 bp and 6851–6950 bp as peaks in (2) and (3) (Fig. 2). In the coding region, 15 and 14 sites in assembly of (2) and (3), respectively, were different from that of (1), possibly due to the homo-polymer errors in Nanopore sequencing. Therefore, we combined the sequence of coding region from assembly of (1) and the CR from assembly of (3) as the final complete nucleotide sequence of *B. japonica* mt-genome (Fig. 1), resulting in a total mt-genome length of 22,274 bp. The mt-genome includes 6,929 bp of CR comprised of conserved sequence block (CSB) 1–3, termination associated sequence (TAS), replication origin of H-strand, and a total of 227 repeat motif sequences (24 motif A repeats and 203 motif B repeats) (Fig. 3).

**Fig. 2.**
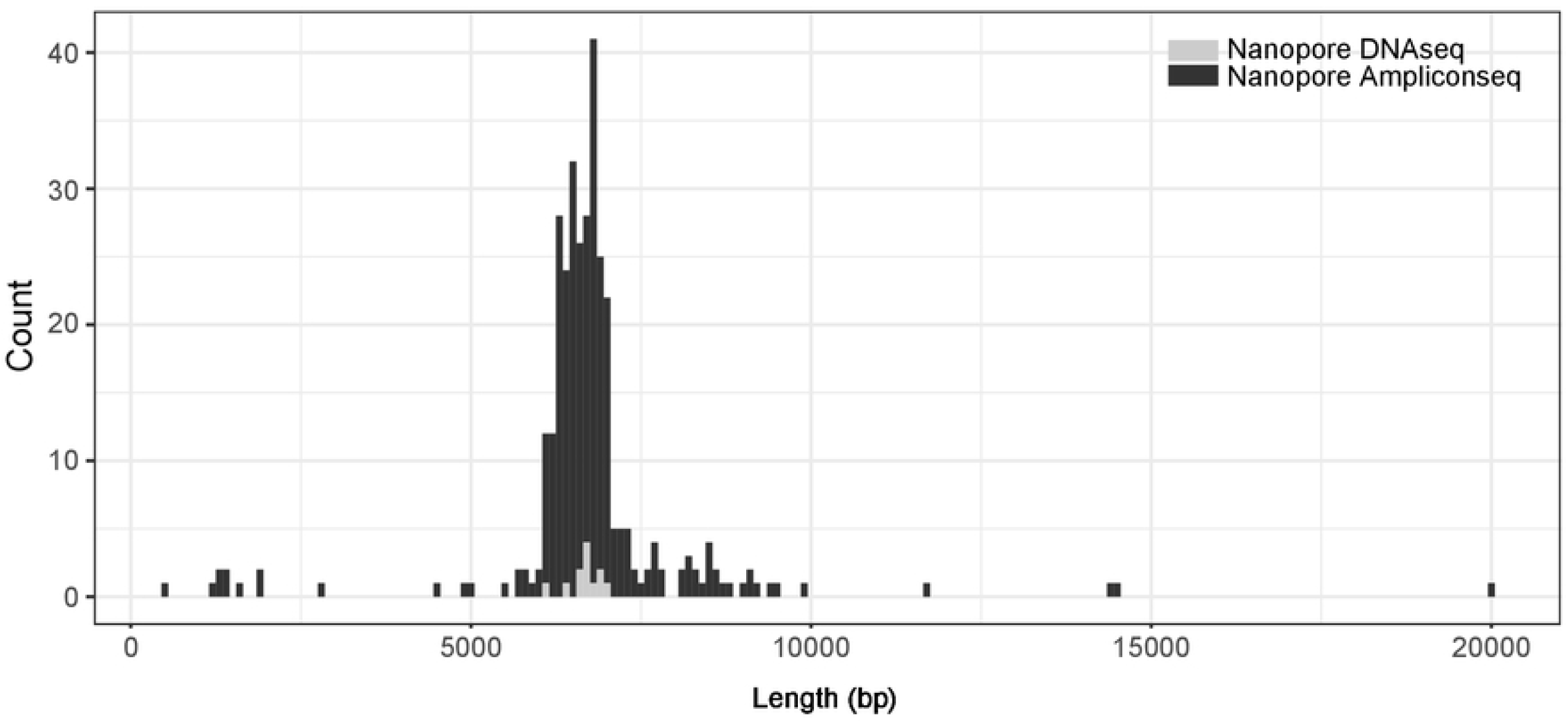
Representative histogram showing length distribution of the raw reads from the control region.

**Fig. 3.**
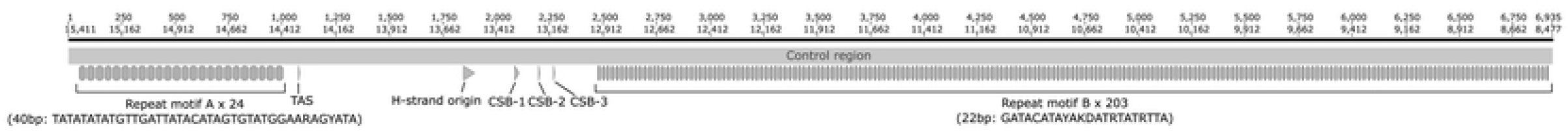
Distribution of repeat motifs in the control region of *B. japonica* mitochondrial genome. The upper and lower numbers indicate nucleotide positions from the 5’ end of the control region and from 5’ end of the ND5 gene, respectively.

In the final complete nucleotide sequence of the *B. japonica* mt-genome, genes for 13 proteins, 2 ribosomal RNAs, and 22 transfer RNAs were annotated. These coding regions started with the ATG codon except COX1 (ATA), ND2 (ATT), ND4L (ATC), ND6 (ATT), and Cytb (ATT). COX1 and ATP8 were terminated by TAA, and COX3, ATP6, ND1, ND3, ND4, and ND4L were terminated by an incomplete stop codon, T. Moreover, COX2, ND5, and Cytb were terminated by AGA, and ND6 and ND2 were terminated by AGG and TAG, respectively. The ND5 gene was located next to the CR. The final assembly was deposited in the DDBJ database under accession number LC739528.

## Discussion

Using short and long read sequencers, we succeeded in the rapid and collective determination of the complete *B. japonica* mt-genome. The present assembly using only short reads (1) results in a short CR (229 bp) with a limited number of repeat motifs. This may be because read length is not enough to reconstruct the whole sequence, and the sequence of repeat region is not defined unless the reads cover both start and end of each repeat region. In addition, the number of repeat motifs is variable in both raw DNA (2) and amplicon (3) (Fig.2); this variability is attributed to somatic mutation (heteroplasmy) and/or PCR error. Since the advent of the Illumina sequencer, mt-genomes of many species have been determined using only short read sequencing with fully automated assemblers (e.g. MitoZ). Although amphibians tend to have a longer CR with repeat motifs, many published mt-genome assemblies of amphibian species may lack CR sequence because they are assembled using short reads only. Therefore, for precise comparison of mt-genomes among species, a careful strategy is necessary to assemble the sequence data.

From the gene annotation of the *B. japonica* mt-genome, we confirm that the resulting gene arrangement is the same as that of the *B. buergeri* mt-genome, indicating that the rearrangement of the ND5 gene may have occurred in the ancestral lineage of genus *Buergeria* after divergence from other lineages of Rhacophorid species. However, the length of the *B. japonica* mt-genome (22,274 bp) is longer than that of *B. buergeri* (19,959 bp), the latter having been accurately determined through sequencing a series of deleted subclones of PCR fragments from *B. buergeri* CR [7]. This difference is due to the presence of much a longer CR in the *B. japonica* (∼7 kbp) compared to *B. buergeri* (∼4.6 kbp). In addition, the two types of repeat motifs observed in the *B. japonica* CR are not homologous with those found in *B. buergeri*; thus, the repeat motifs in *B. japonica* may have accumulated independently over a short period after divergence from the other *Buergeria* species (∼10 – 30 Ma) [8,16].

Intraspecific variation in the length of CR occurs in the eastern spadefoot toad (*Pelobates syriacus*) [32]. Longer mt-genomes are also identified in rain frogs inhabiting sand areas, having adapted to dry and hot environments in East and Southern Africa: *Breviceps adspersus* (28,757 bp), *Breviceps poweri* (28,059 bp), and *Breviceps mossambicus* (22,543 bp) [3]. These previous findings together with our present results suggest that similar evolutionary selective forces acting on mt-genomes, especially in the CR, occur under extreme conditions. Functions of the CR include regulation of mitochondrial gene expression and DNA replication through the formation of D-loop. Recently, R-loops have also been shown to be important for mitochondrial DNA metabolism by the third strand of RNA in the CR [29]. Elucidating the relationship between the tandem repeat motifs in the CR and the mitochondrial gene expression pattern may provide important insights into the thermal adaptation of ectotherms.

## Acknowledgement

We would like to thank Dr. Quintin Lau for English editing. This study was supported by JSPS KAKENHI Grant Number [18K06365, 21K06125] to T.I.

